# Divergent NUMT mimicking functional coding region of the mitochondrial genome

**DOI:** 10.64898/2026.09.18.752603

**Authors:** Marko Prous

**Affiliations:** Ecology and Genetics Research Unit, PO. Box 3000, FI-90014 University of Oulu, Finland; Museum of Natural History, University of Tartu, Vanemuise 46, 51003 Tartu, Estonia

## Abstract

Using long read sequencing, we report here a NUMT (nuclear fragment of mitochondrial origin) in a sawfly species *Euura vittata* (Hymenoptera, Tenthredinidae) that contains the entire coding region with all the genes in the same order as in the mitochondrion. The NUMT is 3.2% different from the mitochondrial sequence (4.1% divergence for COI barcode region), yet appears entirely functional when assuming mitochondrial genetic code. The NUMT is unlikely to be functional because of the usage of standard genetic code in the nuclear genome. Examination of the NUMT containing contig of the whole genome assembly revealed no assembly issues and showed depth of coverage as expected for single copy nuclear genes, strongly suggesting nuclear integration rather than heteroplasmy. Based on mitochondrial COI data, the NUMT is most similar to a cluster of sequences containing mostly *Euura tillbergi* and related species (but also a few *E. vittata* specimens). The NUMT, therefore, may result from recent mitochondrial introgression explaining its divergence and misleading functional appearance. The implications are that reports of divergent heteroplasmy need to be validated with long read or RNA sequencing, or by other appropriate experiments.

## Introduction

Prevalence of mitochondrial DNA insertions in the nuclear genome (NUMTs) varies among taxa [1–4]. Vast majority of NUMTs are short [2,5] because they usually degrade quickly in the nuclear genome where mitochondrial sequences are not functional. Still, complete or near complete mitochondrial genome insertions are known [6–9] and can even be present in multiple (tandem) copies as mega-NUMTs [10,11]. The very long NUMTs are clearly recognizable as such (having stop codons and frameshift indels), even when almost identical (99.9% similarity) to the mitochondrial genome [12]. Recently, Chen et al [4] scanned more than 1000 genomes of avian and mammalian species and found thousands of divergent NUMTs that contain genes with intact mitochondrial reading frames. Also, the recently reported mega-NUMT by Montoliu-Nerin et al. [11] contains 12 copies of complete or near complete mitochondrial genomes, one of which has 9 protein coding genes out of 13 with intact reading frames. However, we are not aware of any study that has reported NUMTs containing the entire mitochondrial coding region which is highly divergent compared to the mitochondrial genome yet shows no signs of pseudogenisation.

Based on mitochondrial COI amplicons (standard DNA barcoding marker for animals), high prevalence of apparently functional intraindividual variants (up to 9% different) has been reported in hundreds of sawfly (Hymenoptera) species [10], even based on ∼1100 bp amplicons [14]. This has led to suggestions that divergent intraindividual mitochondrial variants in sawflies could represent heteroplasmy rather than NUMTs [14,15]. Similar cases of divergent heteroplasmy have been reported in numerous other taxa based on amplicon [16– 18] and / or short read whole genome sequencing [19–21]. Based on the whole genome long read sequencing results reported here, most of these studies reporting divergent heteroplasmy may need to be re-evaluated.

## Material and methods

Three adult sawflies of *Euura vittata* group (Hymenoptera, Tenthredinidae) were selected for genome sequencing because of indications of multiple intraindividual mitochondrial variants in this group based on previous COI amplicon sequencing [14]. The specimens were collected in Finland (table with full specimen data is included as supplementary material). For whole genome sequencing, DNA was extracted with high salt method [22]. The whole body or whole body without the abdomen of the adult specimens were used for extraction. To facilitate release of high molecular weight DNA, the samples were ground to powder (dried ethanol samples or live specimens killed in liquid nitrogen) with sterile pestles before addition of lysis buffer and proteinase K.

Nanopore sequencing libraries were prepared with Ligation Sequencing Kit V14 or Native Barcoding Kit 24 V14 (Oxford Nanopore Technologies, ONT). Raw signal data was basecalled with ONT Dorado 1.0.0-1.2.0 in sup mode (v5.2 basecalling model) and including move tables (‘--emit-moves’ option) for higher accuracy assembly polishing.

For assembly we used Hifiasm 0.25.0 [23] in nanopore mode (--ont), which is suitable for nanopore reads and repetitive genomes. This is critical to recover NUMTs similar to mitochondrial genome. Several assemblies with different read length cutoffs (500–10000 bp) were performed to find optimal assembly based on BUSCO results. The whole genome assembly was polished one time with dorado polish (model dna_r10.4.1_e8.2_400bps_sup@v5.2.0_polish_rl_mv). The potential mitochondrial (mt) contigs were identified with blastn [24] using cox1 sequences of the same species as query. For high depth organellar genomes, hifiasm often creates tandem repeats of them. We used SeqKit [25] to set the start of trnaI gene as the start position and then separate the individual pseudo-repeats. To identify the trnI and the other mitochondrial genes, we used MITOS [26] implemented in usegalaxy.eu [27]. We compared the individual mitochondrial genome pseudo-repeats of the assembly to confirm they were identical or near identical (differences typically only in the number of AT repeats or lengths of homopolymers) at least in the coding region. The long (longer than the coding region) and highly repetitive non-coding control region varied between the complete mitochondrial genome pseudo-repeats due to differences in number of various subrepeats. We selected one of the middle size mitochondrial genome assembly repeats for validation. First, we used SeqKit to subsample reads so that mitochondrial genome depth of coverage was roughly between 100-1000X (min read length 4 kb - 15 kb, max about the length of mt genome) to reduce nuclear genome coverage. To further limit mapping of NUMT containing reads to the mitochondrial genome assembly, we used minimap2 [28] with the preset -x asm5 that enables mapping only the high similarity reads (∼0.1% sequence divergence). We then visually inspected the mapping with Tablet [29] to check if depth of coverage was similar (e.g. around 100–200X) along the assembly and to correct small errors based on the mapped reads, typically single nucleotide indels next to or in the middle of long dinucleotide AT repeats: 0–1 substitutions (0 in the coding region), 2–97 single nucleotide indels (0–1 in the coding region) depending on the assembly.

To assess the genome assembly quality based on BUSCO score [30], we used Hymenoptera protein set (hymenoptera_odb12) with *compleasm* [31].

Probably due to high sequencing depth of mitochondrial genome, variation in the highly repetitive non-coding control region (differences in the number of repeats), sequencing (e.g. homopolymers, strand bias) and other errors (chimeric reads), hifiasm assembly produced several mitochondrial contigs (7–218 contigs depending on the assembly), most of which were short (<10 kb). These contigs were identified with blastn (using the validated mitochondrial genome as query) and the read support for these contigs was checked based on the mapping (minimap2 -x asm5) of the genome assembly. The contigs that could entirely be considered as fragments of the mitochondrial genome (closer to the mitochondrial genome than to the other contigs along the entire length) or were partly mitochondrial and partly unsupported by original reads (assembly errors) were removed from the final assembly and replaced with the validated mitochondrial genome.

To confirm the validity of the NUMT containing region, the reads were mapped (minimap2 preset -x asm5) to the assembly and checked with Tablet and IGV [32]. To identify possibly NUMT junctions, the NUMT containing region was aligned to the closest mitochondrial genome sequence using MAFFT v7.526 [33].

GC content along contigs was calculated for 1000 bp windows sliding by 500 bp with bedtools (https://bedtools.readthedocs.io/) (commands *bedtools makewindows* and *bedtools nuc*). Sequencing depth for contig positions was obtained with *samtools depth* [34]. Barplots of GC content and depth were made in R.

The p-distances (proportion of nucleotide differences) and non-synonymous (amino acid-changing, dN) to synonymous (silent, dS) substitution ratios were calculated in R with the package *ape* [35].

To place the three species for which the whole genomes were sequenced here (the only genomes available so far for *Euura vittata* group) in the phylogenetic context, we used the genes widely sequenced for sawflies (see Prous et al. [11]): cytochrome c oxidase subunit I (COI), 6-phosphogluconate dehydrogenase (PGD), triose-phosphate isomerase (TPI), F2 copy of elongation factor 1_α_ (EF1a F2), sodium/potassium-transporting ATPase subunit alpha (NaK), h1 copy of heat shock protein 83 (HSP90 h1), DNA dependent RNA polymerase II subunit RPB1 (POL2), and transformation/transcription domain-associated protein (TRRAP). Most of these sequences used here were published before [14], the newly generated ones (besides the sequenced genomes) were amplified and sequenced as described in Prous et al. [14].

For phylogenetic analyses (separately for COI and nuclear genes) of nucleotide alignments (amino acid alignments would be uninformative due to low or even zero divergence), we used IQ-TREE ver. 3.0.1 [36]. By default, IQ-TREE runs ModelFinder [37] to find the best-fit substitution model and then reconstruct the tree using the model selected according to Bayesian information criterion (BIC). We complemented this default option with a SH-like approximate likelihood ratio (SH-aLRT) test [38], ultrafast bootstrap [39] with 1000 replicates, and in case of concatenated nuclear genes, gene concordance factor [40] to estimate robustness of reconstructed splits.

The basecalled reads used here for genome assemblies have been submitted to NCBI trace archive (accessions SRR40721943–SRR40721945). The newly obtained genome assemblies (accessions JCFLQJ000000000-JCFLQL000000000 under BioProject PRJNA1525494) and amplicon sequences (accessions QB012738, QB020663–QB020667) have been submitted to GenBank.

## Results

BUSCO scores of the polished assemblies of three *Euura vittata* group specimens were 96.62–98.72% (Table 1). Nuclear genome assembly sizes were 300–317 Mb and mitochondrial genomes 48–55 kb (Table 1). In *Euura vittata*, a NUMT representing the entire section of the coding region flanked on both sides by large fraction of the non-coding control region was detected. For simplicity, we use “coding region” to refer to the region containing all the mitochondrial genes (13 protein coding genes, 22 tRNA and 2 rRNA genes). The NUMT was located within a 4.5 Mb contig, detectable also based on low GC content (Fig. 1). Based on the comparison with the mitochondrion of *E. minivittata* (the closest mitochondrial sequence available to the NUMT), the NUMT was 30 247 bp long. Due to high variability of the non-coding control region of the mitochondrion, the estimated length might not be exact and complete integration of the control region (in addition to the complete coding region) cannot be excluded (even if the length is significantly shorter than the mitochondria sequenced here). 4883 bp downstream of the 30 kb NUMT, there was a separate 2670 bp section representing part of the control region (Fig. 1). Considering only reads extending at least 1000 bp to both sides of the nuclear-NUMT junctions, nine (start) and eight (end) supported the 30 kb NUMT localisation in the 4.5 Mb contig (15 unique reads, two of which span the 30 kb region entirely) (Figs S1–S2). For the 2670 bp NUMT fragment, nine (start) and five (end) reads of at least 2 kb fully supported the localisation (12 unique reads, two of which span the 2670 bp region entirely) (Figs S3–S4). The coding region (15 196 bp) of the NUMT is 3.2% different (Table 2) to the coding region (15 239 bp) of the mitochondrial genome of the same specimen and all the 37 mitochondrial genes are in the same order between them. Remarkably, all the genes of this NUMT appear entirely functional when assuming mitochondrial genetic code. Additionally, non-synonymous to synonymous substitution ratio (dN/dS) calculations indicate that the NUMT sequence has been under strong purifying selection with dN/dS below 0.1 (Table 3), supporting its recent origin from true mitochondrion. No such NUMTs were detected in closely related species of *E. hedstroemi* or *E. minivittata*. Since the nuclear genome uses standard genetic code, the NUMT is unlikely to be functional.

**TABLE 1.** Genome assembly statistics of *Euura vittata* group specimens.

| **Sample** | ***E. vittata* ZMUO.079758 female** | ***E. hedstroemi* ZMUO.067447 male** | ***E. minivittata* ZMUO.067456 male** |
|---|---|---|---|
| **Assembly length** | 316.6 Mb | 302.5 Mb | 300.3 Mb |
| **Contigs** | 987 | 554 | 56 |
| **N50** | 2.8 Mb | 3.8 Mb | 14.1 Mb |
| **max** | 9.5 Mb | 10.9 Mb | 43.5 Mb |
| **min** | 1436 bp | 664 bp | 8396 bp |
| **Depth** | 31.0 | 20.4 | 54.6 |
| **mtDNA length** | 48.3 kb | 54.3 kb | 55.3 kb |
| **mtDNA depth** | 2397 | 1382 | 6099 |
| **BUSCO single + duplicated** | 98.37% | 99.3% | 99.3% |
| **single** | 96.62%, 5719 | 98.61%, 5837 | 98.72%, 5843 |
| **duplicated** | 1.76%, 104 | 0.66%, 39 | 0.63%, 37 |
| **fragmented** | 0.35%, 21 | 0.15%, 9 | 0.15%, 9 |
| **missing** | 1.27%, 75 | 0.57%, 34 | 0.51%, 30 |

**TABLE 2.** p-distances among the mitochondrial genomes and the NUMT.

|  | ***E. vittata*** | ***E. hedstroemi*** | ***E. vittata* NUMT** |
|---|---|---|---|
| ***E. hedstroemi*** | 0.0123 |  |  |
| ***E. vittata* NUMT** | 0.0325 | 0.0341 |  |
| ***E. minivittata*** | 0.0319 | 0.0338 | 0.0156 |

**TABLE 3.** Non-synonymous to synonymous substitution ratios (dN/dS) among the mitochondrial genomes and the NUMT.

|  | ***E. vittata*** | ***E. hedstroemi*** | ***E. vittata* NUMT** |
|---|---|---|---|
| ***E. hedstroemi*** | 0.0881 |  |  |
| ***E. vittata* NUMT** | 0.0625 | 0.0649 |  |
| ***E. minivittata*** | 0.0687 | 0.0741 | 0.0671 |

**Figure 1.**
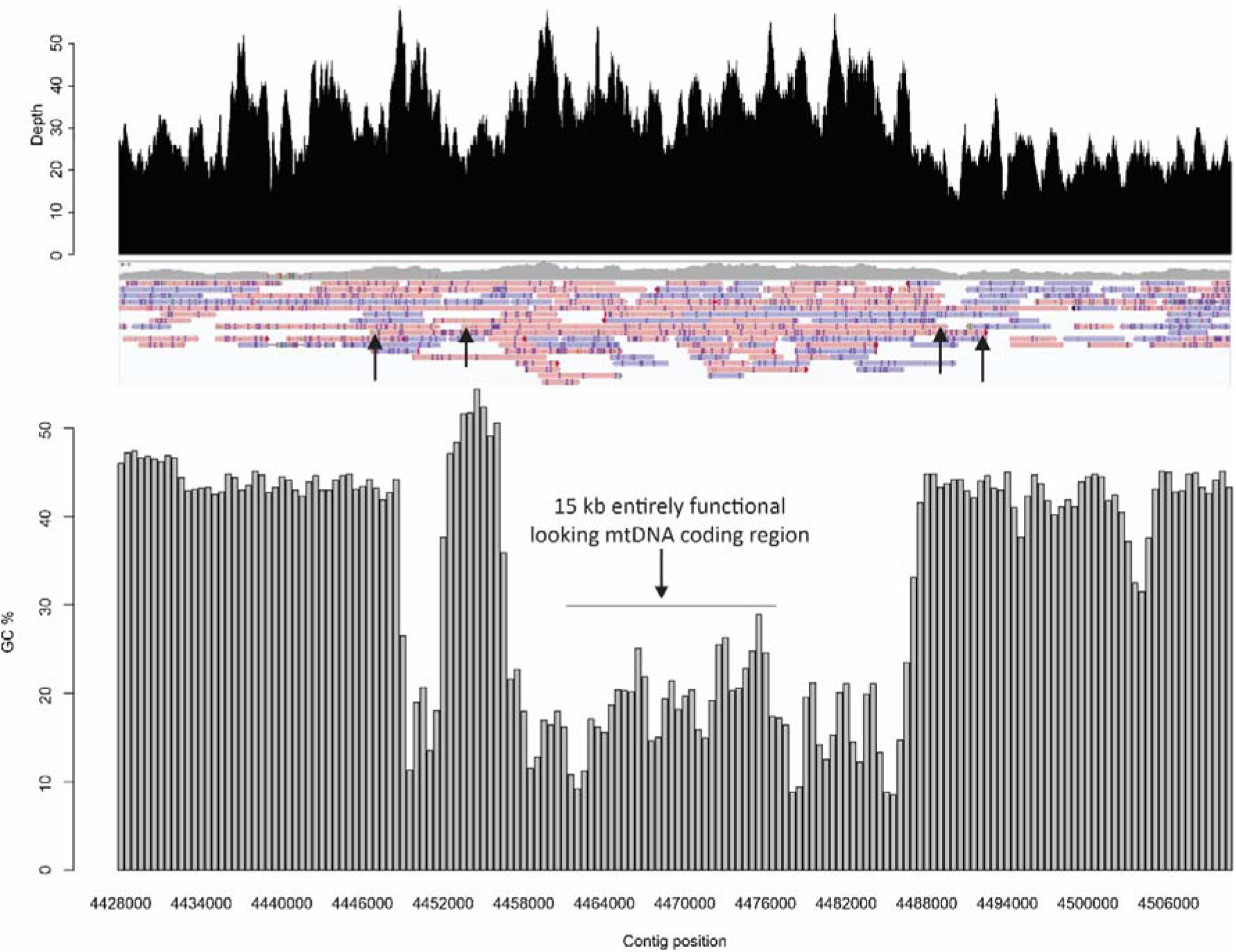
Sequencing depth based on at least 200 bp reads (top), mapped reads of at least 2 kb (middle), and GC content (%, bottom) along the NUMT containing 4.5 Mb contig of *Euura vittata*. Average sequencing depth for the entire contig is 32.2X, for the contiguous 30 kb NUMT containing region (including the coding region indicated in the figure) 37.5X. Sequencing depth for the mitochondrial DNA is 2397X. Arrows in the middle panel indicate start and end positions of two reads that span the entire 30 kb NUMT region, one of them extending also beyond the shorter (2670 bp) NUMT fragment with low GC content on the left.

The NUMT is closely related to the mitochondrial genome of *Euura minivittata* with divergence of 1.6% (Table 2). To put the NUMT in more detailed phylogenetic context, we used ∼1000 bp fragment of COI [14] to build a maximum likelihood tree for *Euura vittata* group, including all detected intraindividual variants that appear functional. Most species are not monophyletic based on COI (Fig. 2), which is a common pattern in sawflies [14,41]. In contrast, monophyly is well supported for most species based on nuclear genes (Fig. 3). The NUMT of *Euura vittata* specimen ZMUO.079758 is identical to several COI secondary variants (less frequent based on PCR sequencing output than the dominant variant) of some other *E. vittata* specimens (Fig. 2), which likely also represent NUMTs. The next closest COI sequences to the NUMT (Fig. 2) come from *E. vittata* (p-distance 0.5–0.6%; most of which are secondary variants), *E. tillbergi* (0.7–1.4%), and *E. krausi* (1.0–1.3%).

**Figure 2.**
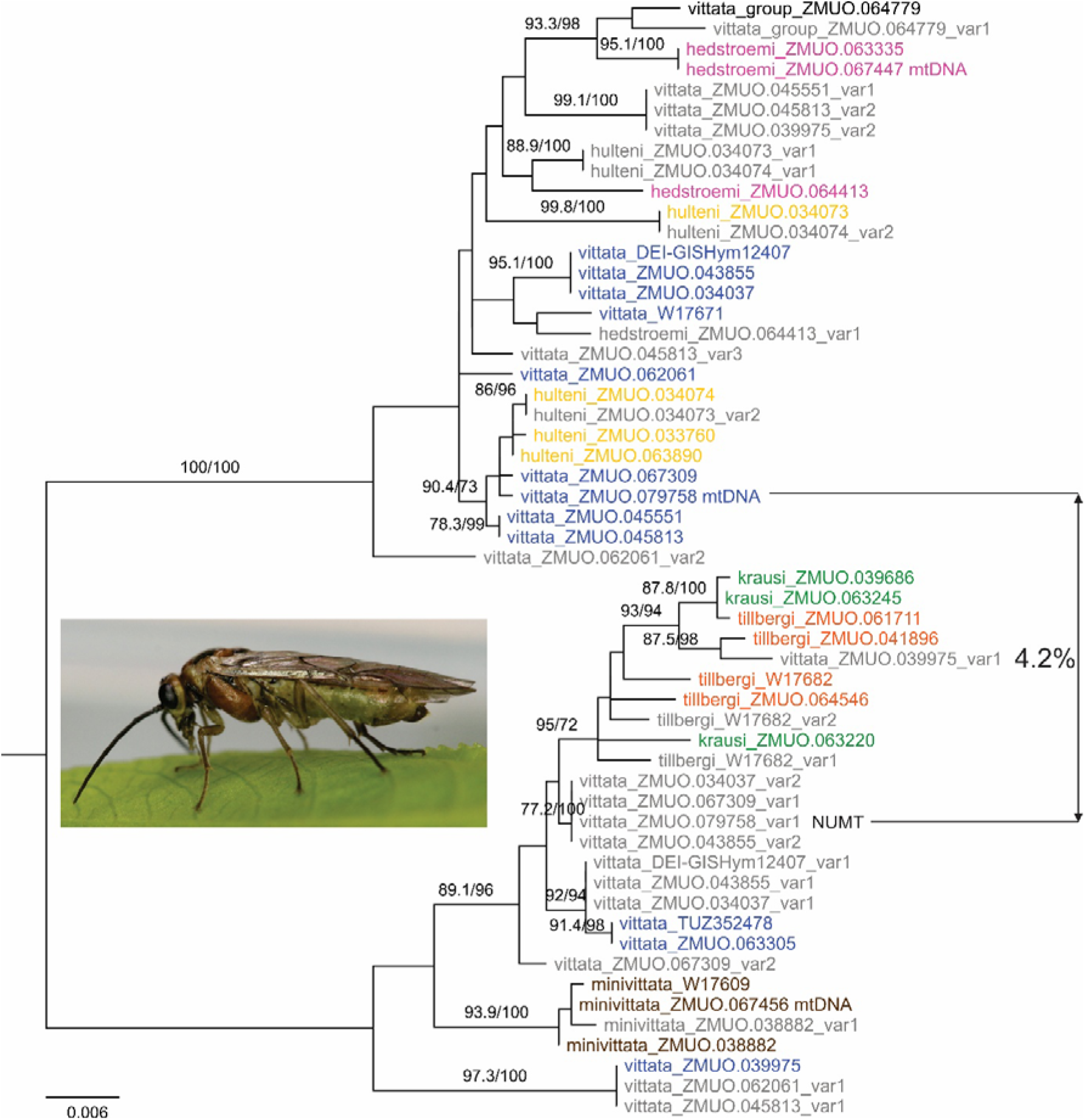
Maximum likelihood tree (TN+F+I model) of COI sequences (1078 bp) from 30 specimens (55 variants) of *Euura vittata* group. The sequences marked with “mtDNA” are confirmed mitochondrial sequences based on the whole genomes sequenced here. The “NUMT” is part of the NUMT of *Euura vittata* ZMUO.079758. All the other sequences are based on PCR and those in grey colour are secondary variants (less common based on sequencing output). Sequence divergence between mitochondrial COI fragment and the NUMT fragment of ZMUO.079758 is shown. The photo shown is of *Euura vittata* TUZ615924. Numbers at branches show SH-aLRT support (%) / ultrafast bootstrap support (%) values. Values of only well supported branches (>90 for at least one of the support values) are shown.

**Figure 3.**
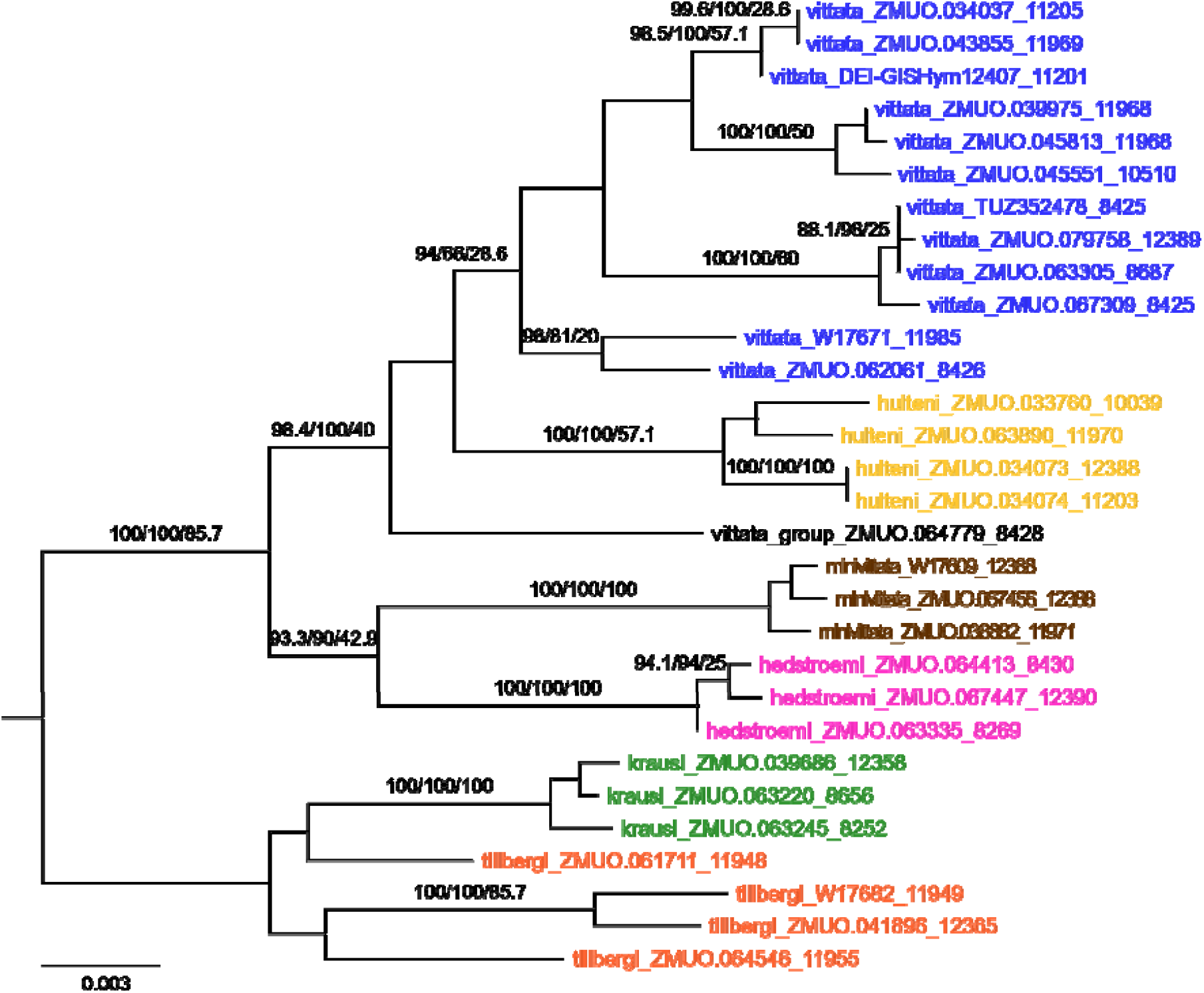
Maximum likelihood tree (TIM+F+I+G4) based on concatenation of 7 nuclear genes (alignment length 12424 bp) of the same 30 specimens as in Fig. 1. Numbers at branches show SH-aLRT support (%) / ultrafast bootstrap support (%) / gene concordance factor (%) values. Values of only well supported branches (>90 for at least SH-aLRT or ultrafast bootstrap) are shown. The numbers at the end of the tip labels refer to the sequence length available.

## Discussion

NUMTs have been known for long time and have been studied extensively. Vast majority of NUMTs are short [1–3,5], but those representing large fraction or complete mitochondrial genome, even in multiple copies, are known [6–12]. These NUMTs have been recognizable as such due to stop codons and / or frameshift indels (insertions or deletions), though NUMT diagnostic mutations might naturally be absent in very young insertions (identical or very similar to the species’ mitochondrial genome). Nevertheless, divergent NUMTs of individual genes with intact mitochondrial (mt) reading frames (assuming mitochondrial genetic code) have been found [4]. In some NUMTs representing (near) complete mt genome, some or most of the genes can be with intact mitochondrial frames [9,11]. Remarkably, as we have shown here, even divergent (3.2% different to the mtDNA) complete coding region of the mitochondrial genome can be present in the nuclear genome without any NUMT diagnostic mutations. How is it possible that of all the mutations (roughly 400 mutations in case of 3.9% divergence for protein coding sequences), none have introduced stop codons or frame shifts when assuming mitochondrial genetic code? This could be explained by recent enough mitochondrial introgression from a different species [4]. For example, *Trachypithecus* monkey species (*T. pileatus* and *T. shortridgei*) have a NUMT covering 92% of the mt genome (7 protein sequences out of 13 of this NUMT have premature stop codons). However, the NUMT is closer (divergence 5.8–6.5%) to *Semnopithecus* than to the other *Trachypithecus* (divergence 14.4–15.1%) [9]. Similarly, the NUMT in *Euura vittata* described here belongs to a clade mostly represented by *E. tillbergi* and *E. krausi* (Fig. 2). However, the closest possibly non-NUMT sequences are a few sequences of *E. vittata* (Fig. 2). Mitonuclear discordance (phylogenetic conflict between nuclear and mitochondrial genes) is common in sawflies and is most likely resulting from mitochondrial introgression [14,41,42]. Within *Euura vittata* group, *E. tillbergi* and *E. krausi* form a separate subgroup (*tillbergi* subgroup) based on nuclear genes (Fig. 3) and morphology [14]. This suggests introgression of mtDNA from *tillbergi* subgroup to *E. vittata*, explaining the origin of the NUMT, its divergence to the mtDNA, and its apparent functionality when assuming mitochondrial genetic code.

In light of the results reported here, previous mitochondrial COI amplicon data (∼650 bp or even ∼1100 bp) in sawflies [13,14] might rather suggest high prevalence of long NUMTs (at least large sections of the COI gene) with intact mitochondrial reading frames than presence of heteroplasmy. Divergent heteroplasmy is increasingly reported, but most of these reports would need confirmation based on our results. For example, Kastally & Mardulyn [19] reported in a beetle species the presence of two divergent mitochondria (>1% divergence) within individuals based on Illumina short read sequencing and population screening of PCR amplicons using primers specific to the two mitochondrial variants. Meza-Lázaro et al. [20] also reported divergent heteroplasmy (up to 5.8% divergence) based on whole genome Illumina sequencing in a species complex of *Ectatomma* ants, which interestingly seem to be hybridizing. These and similar reports of heteroplasmy prompted Allison et al. [43] to employ population genetic modelling which indeed suggested that biparental inheritance of mitochondria is favoured under moderate levels of gene flow in hybridizing populations.

Furthermore, failure to eliminate paternal mtDNA has been suggested to be more likely with increasing genetic distance between the parents [17,44]. Therefore, empirical and theoretical research seem to agree that divergent heteroplasmy might not be unusual. Since NUMT integration could be common, continuous, and highly dynamic, at least in some taxa, much of the previous empirical research on divergent heteroplasmy might need re-evaluation based on long read sequencing or other appropriate experiments. Nevertheless, divergent heteroplasmy cannot be entirely excluded (besides taxa like some Bivalvia with doubly uniparental inheritance of mitochondria), but more evidence besides Illumina sequencing or PCR based population screening should be presented before reporting heteroplasmy. Functional appearance of mitochondrial sequences is insufficient evidence to assume its localisation in the mitochondrion.

## Data accessibility

Raw sequences have been submitted to NCBI SRA. Genome assemblies and amplicon sequences have been submitted to GenBank. Alignment files (COI and concatenated nuclear genes) used to construct maximum likelihood trees, corresponding specimen data, and additional supporting text and figures (S1–S4) are included as electronic supplementary material.

## Acknowledgements

I would like to thank Marko Mutanen (University of Oulu) for the sawfly samples and generous support throughout the project. Suggestions and corrections by anonymous referees helped to improve the manuscript. This research received support from the Estonian Research Council grant STP42 as well as from the Biodiverse Anthropocenes research profiling programme of the University of Oulu. MP is currently supported by the Kvantum Institute of the University of Oulu project “Developing a genomic blueprint for a bio-literate future”. Some of the analyses (basecalling, genome polishing) were run using high performance computing (HPC) services at CSC (https://research.csc.fi/) or HPC center of University of Tartu (https://hpc.ut.ee/).

