## Supporting Information for "Divergent NUMT mimicking functional coding region of the mitochondrial genome"

Marko Prous

**This file includes:**

**Supporting text**

**Figures S1–S4**

**Other supporting materials for this manuscript include:**

**Datasets S1**

Fasta alignment (cox1.fas) of COI sequences (1078 bp) used to construct ML tree in Fig. 2.

**Datasets S2**

Fasta alignment (nuclear_concat7genes.fas) of concatenated 7 nuclear genes (12424 bp) used to construct ML tree in Fig. 3

**Datasets S3**

Excel table (specimen_data.xlsx) containing occurence data and GenBank accessions for the specimens used in the phylogenetic analyses.

**Supporting text**

**Commands for genome assembly, polishing, mapping etc.**

Example of hifiasm command with read length cutoff 1000 bp:

hifiasm --ont --rl-cut 1000 -o assembly -t 12 reads.fastq

Converting hifiasm assembly graph output to fasta file:

awk '/^S/{print ">"$2"\n"$3}' assembly.bp.p_ctg.gfa | fold > assembly.fasta

Downloading Hymenoptera protein set for compleasm:

compleasm.py download hymenoptera_odb12 -L buscolin

Running compleasm:

compleasm_kit/compleasm.py run -t8 -l hymenoptera -L buscolin -a assembly.fasta -o Output

Dorado polishing:

rm *.bai # removing index files if present from previous analyses

rm *.fai

samtools faidx assembly.fasta

dorado aligner assembly.fasta reads.fastq.gz | samtools sort > alignment.bam

samtools index alignment.bam

dorado polish alignment.bam assembly.fasta --ignore-read-groups > polished_assembly.fasta

"--ignore-read-groups" was used to avoid dorado polish fail in cases when reads are coming from more than one sequencing run.

Example of assembly mapping:

minimap2 -a --sam-hit-only --secondary=no -x asm5 -t 8 assembly.fasta reads.fastq | samtools sort > alignment.bam

samtools index alignment.bam

Average sequencing depth:

samtools depth alignment.bam | awk '{sum+=$3} END {print "Average = ",sum/NR}'

Sequencing depth for specific contig (e.g. ptg000044l) positions (e.g. 4427000-4510500) and plotting in R:

samtools depth -r ptg000044l:4427000-4510500 alignment.bam > depth.txt

genome = read.table("depth.txt", sep = "\t", quote = "", header = F) # reading table in R

barplot(genome$V3, names.arg = genome$V2, xlab = "Contig position", ylab = "Depth")

Example of seqkit command to filter reads based on the length:

seqkit seq -m 2000 -M 50000 reads.fastq > output.fastq

Setting standard start position (trnI gene) for mitochondrial genome assembly:

trnI=AATGAAGTGCCTGAATAAAAGGAT # example of start sequence of trnI gene

seqkit locate -p $trnI mt_contig.fasta | tail -n+2 > start.txt

seqkit restart -i $(cut -f5 start.txt) mt_contig.fasta > output.fasta

Plotting GC content:

seqkit grep -p ptg000044l polished_assembly.fasta > input.fas # finding contig ptg000044l from the assembly

echo -e $(seqkit seq -n input.fas)'\t'$(seqkit stats -T input.fas | cut -f5 | tail -n1) > contig.txt # table for contig names and its lengths

rm *.fai

bedtools makewindows -g contig.txt -w 1000 -s 500 > windows.bed

bedtools nuc -fi input.fas -bed windows.bed | sed s/^#// > table.txt

genome = read.table("table.txt", sep = "\t", quote = "", header = T) # reading table in R

barplot(genome$X5_pct_gc[8855:9020], names.arg = genome$X3_usercol[8855:9020], xlab = "Contig position", ylab = "GC %", main = "GC content along the contig containing the NUMT")

**Distance and dN/dS calculations with R package *ape***

alignment = read.FASTA("alignment.fasta")

alignment = as.matrix(alignment)

dist.dna(alignment, model = "raw", pairwise.deletion = T) # p-distances

dnds(alignment, code = 5, codonstart = 1, quiet = T) # dN/dS, only for protein coding sequences, "code = 5" refers to mitochondrial genetic code (5)

**Genome assembly polishing with dorado**

Dorado does not provide detailed output on positions which were changed. In practice, the main changes are in homopolymeres which become longer compared to the unpolished assembly. Accordingly, the polished assemblies of the three *Euura vittata* group specimens were 83328–116956 bp (0.03–0.04%) longer. BUSCO scores remained the same or improved slightly after the polishing. For *Euura hedstroemi* and *E. minivittata*, single copy gene score did not change at all after polishing. For *Euura vittata*, single copy gene score increased from 96.52% to 96.62% (one fewer missing gene and five fewer duplicated).

**Reads crossing NUMT junctions**

Reads at least 2000 bp were mapped to the assembly with minimap2:

minimap2 -a --sam-hit-only --secondary=no -x asm5 assembly.fasta reads2kb.fastq -t 10 | samtools sort -@ 10 > alignment.bam

samtools index alignment.bam

The alignment bam file was visualized with IGV and Tablet. The 4.5 Mb contig (ptg000044l) contained two NUMT fragments interrupted by 4883 bp of nuclear sequence (no homology to mitochondrion): 2670 bp fragment from non-coding control region of the mitochondrion and 30 247 bp fragment containing complete coding region (15 196 bp) in the middle and flanked by fragments of the non-coding control region.

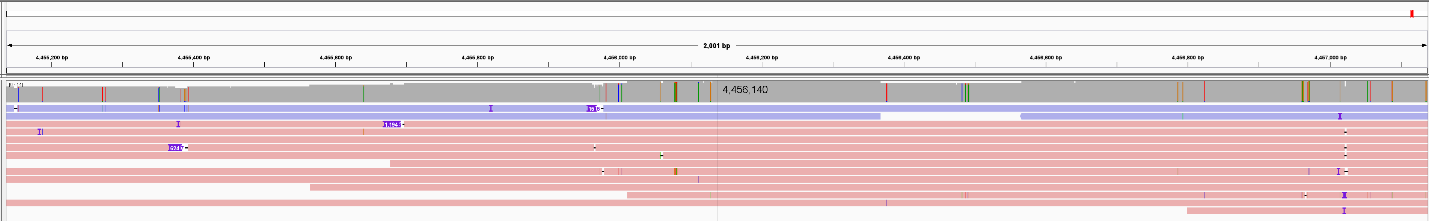

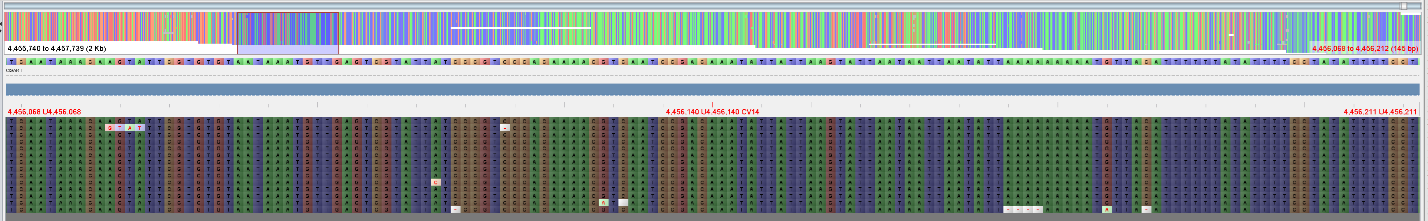

**Figure S1. Top**: 2 kb window around the start (4 456 140) of the 30 kb NUMT fragment visualized with IGV. Reads in blue are reverse complements relative to the contig. Nine reads extend at least 1000 bp to both sides of the junction. **Bottom**: Detailed view with Tablet. 144 bp window highlighted on the top panel is shown in detail below.

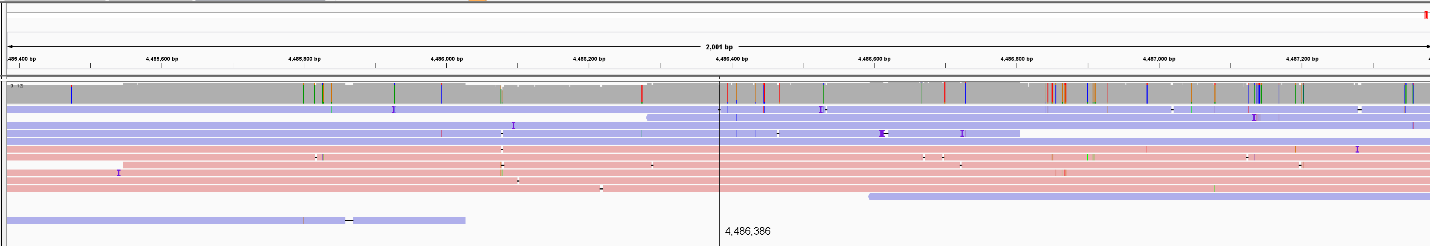

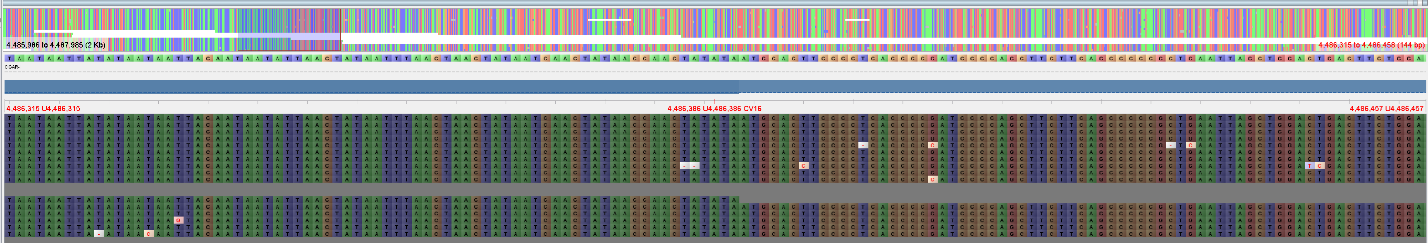

**Figure S2. Top**: 2 kb window around the end (4 486 386) of the 30 kb NUMT fragment visualized with IGV. Reads in blue are reverse complements relative to the contig. Eight reads extend at least 1000 bp to both sides of the junction. **Bottom**: Detailed view with Tablet. 144 bp window highlighted on the top panel is shown in detail below.

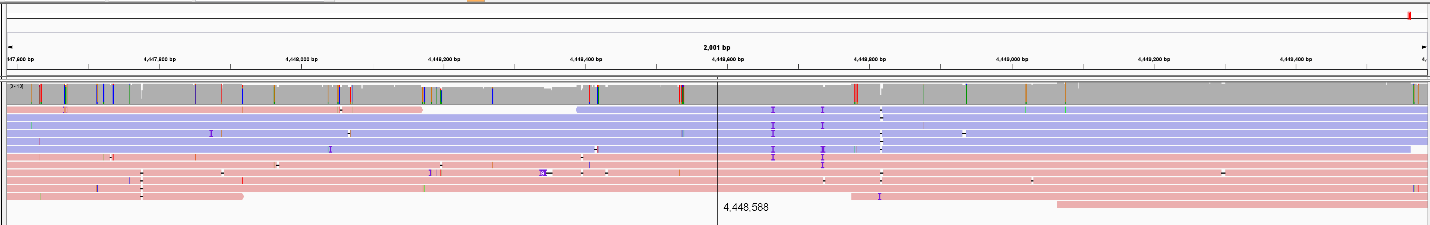

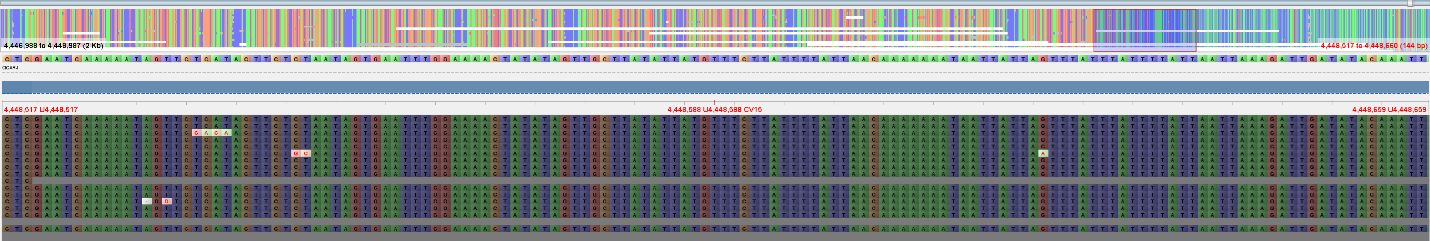

**Figure S3. Top**: 2 kb window around the start (4 448 588) of the 2670 bp NUMT fragment visualized with IGV. Reads in blue are reverse complements relative to the contig. Nine reads extend at least 1000 bp to both sides of the junction. **Bottom**: Detailed view with Tablet. 144 bp window highlighted on the top panel is shown in detail below.

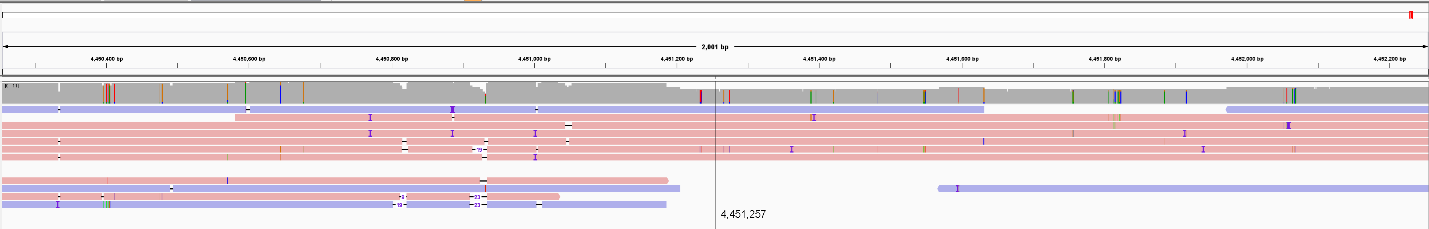

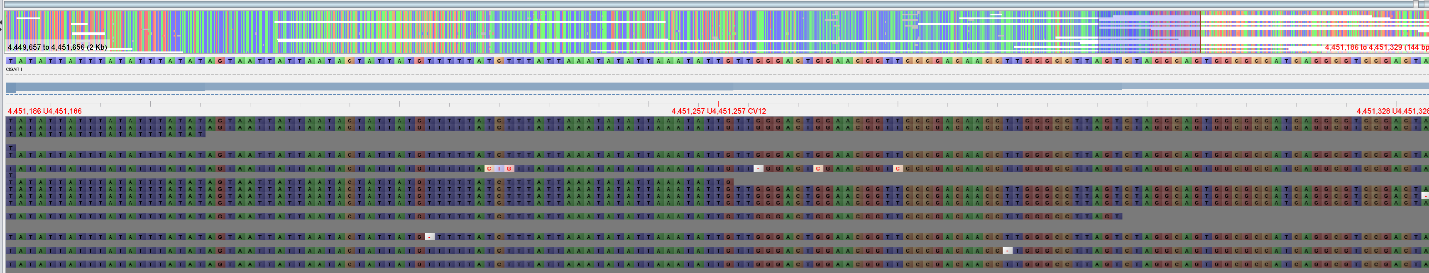

**Figure S4. Top**: 2 kb window around the end (4 451 257) of the 2670 bp NUMT fragment visualized with IGV. Reads in blue are reverse complements relative to the contig. Five reads extend at least 1000 bp to both sides of the junction. **Bottom**: Detailed view with Tablet. 144 bp window highlighted on the top panel is shown in detail below.
